# S100A8/A9 mediate the reprograming of normal mammary epithelial cells induced by dynamic cell-cell interactions with adjacent breast cancer cells

**DOI:** 10.1101/312884

**Authors:** Seol Hwa Jo, Woo Hang Heo, Mingji Quan, Bok Sil Hong, Ju Hee Kim, Han-Byoel Lee, Wonshik Han, Yeonju Park, Dong-Sup Lee, Nam Hoon Kwon, Min Chul Park, Jeesoo Chae, Jong-Il Kim, Dong-Young Noh, Hyeong-Gon Moon

**Author notes:** Address correspondence to: Hyeong-Gon Moon, Seoul National University College of Medicine, 103 Daehak-ro, Jongno-gu, Seoul, 03080, South Korea.

## Abstract

To understand the potential effects of cancer cells on surrounding normal mammary epithelial cells, we performed direct co-culture of non-tumorigenic mammary epithelial MCF10A cells and various breast cancer cells. Firstly, we observed dynamic cell-cell interactions between the MCF10A cells and breast cancer cells including lamellipodia or nanotube-like contacts and transfer of extracellular vesicles. Co-cultured MCF10A cells exhibited features of epithelial-mesenchymal transition, and showed increased capacity of cell proliferation, migration, colony formation, and 3-dimensional sphere formation. Transcriptome analysis and phosphor-protein array suggested that several cancer-related pathways are significantly dysregulated in MCF10A cells after the direct co-culture with breast cancer cells. S100A8 and S100A9 showed distinct up-regulation in the co-cultured MCF10A cells and their microenvironmental upregulation was also observed in the orthotropic xenograft of syngeneic mouse mammary tumors. When S100A8/A9 overexpression was induced in MCF10A cells, the cells showed phenotypic features of directly co-cultured MCF10A cells in terms of *in vitro* cell behaviors and signaling activities suggesting a S100A8/A9-mediated transition program in non-tumorigenic epithelial cells. This study suggests the possibility of dynamic cell-cell interactions between non-tumorigenic mammary epithelial cells and breast cancer cells that could lead to a substantial transition in molecular and functional characteristics of mammary epithelial cells.

## Introduction

In solid tumors, the complex tumor microenvironment controls all steps of tumor progression and metastasis.^1,2^ The tumor microenvironment is comprised of various endogenous and recruited cells that undergo dynamic cell-cell interactions with malignant epithelial cells and contribute to the tumor cell’s behaviors.^3,4^ For example, cancer-associated fibroblasts actively remodel extracellular matrix and immune microenvironment, and cancer-associated adipocytes provide inflammatory milieu that support tumor growth.^5,6^ Moreover, recent efforts to target the immune microenvironment have shown promising therapeutic responses in selected solid tumors.^7^ Therefore, understanding the molecular mechanisms of the tumor-microenvironment interactions can provide scientific basis for developing novel therapeutic strategies that target the tumor microenvironment.^3,8,9^

Normal epithelial cells are closest neighbors to the malignant transformed cells in human epithelial tumors arising from solid organs. During the early steps of carcinogenesis, the normal epithelial cells may exert tumor-suppressive effects by promoting protrusion of transformed epithelial cells from the epithelial layers.^10-12^ However, the tumor-suppressive effects of normal epithelial cells may not last throughout the solid tumor progression. While the normal myoepithelial cells obtained from healthy human breast tissues contribute to the maintaining polarity of mammary epithelial cells and suppress aberrant growth, the myoepithelial cells derived from breast cancer tissues failed to restore physiologic polarity in mammary epithelial cells and showed increased expression of various chemokines such as CXCL12.^13,14^ These reports suggest a potential functional transition of normal epithelial cells caused by adjacent malignant epithelial cells which may contribute the progression of solid tumors.

In this study, we show that breast cancer cells and non-tumorigenic mammary epithelial cells undergo dynamic cell-cell interactions that lead to a substantial reprograming of molecular characteristics of the mammary epithelial cells. The reprograming of normal mammary epithelial cells includes phenotypes changes as well as dysregulations of mRNA expression and cell signaling activities. Our data suggests that S100A8/A9 upregulation in non-tumorigenic mammary epithelial cells may play a critical role in the phenotype shifting induced by adjacent cancer cells.

## Results

### Dynamic interaction between breast cancer cells and non-transformed mammary epithelial cells

First, we determined the presence and the extent of cell-cell interactions *in vitro* between the breast cancer cells and mammary epithelial cells. We co-cultured the RFP-transfected breast cancer cells (MDA-MB-231) with GFP-transfected non-transformed mammary epithelial cells (MCF10A) using in vitro direct co-culture method. While the majority of MCF10A cells maintained the clusters of adherent cells, MDA-MB-231 cells showed spreading patterns of cell growth and the cells infiltrated between the MCF10A cell clusters (Supplementary Figure S1). The time-lapse imaging of the cells showed that MDA-MB-231 cells had more frequent cell movements than the MCF10A cells and the cells showed various dynamic cell-cell interaction patterns (Supplementary Video S1 and S2). MDA-MB-231 cells formed both lamellipodia-like structures for adjacent cells and nanotube-like projections for long-range cell-cell interactions (Figure 1a and 1b, Supplementary Video S1).^15,16^ The lamellipodia-like structures of MDA-MB-231 cells actively contacted the MCF10A cells and a portion of extended lamellipodia could remain as extracellular vesicles which were then engulfed by adjacent MCF10A cells (Figure 1c, Supplementary Video S2).

**Figure 1.**
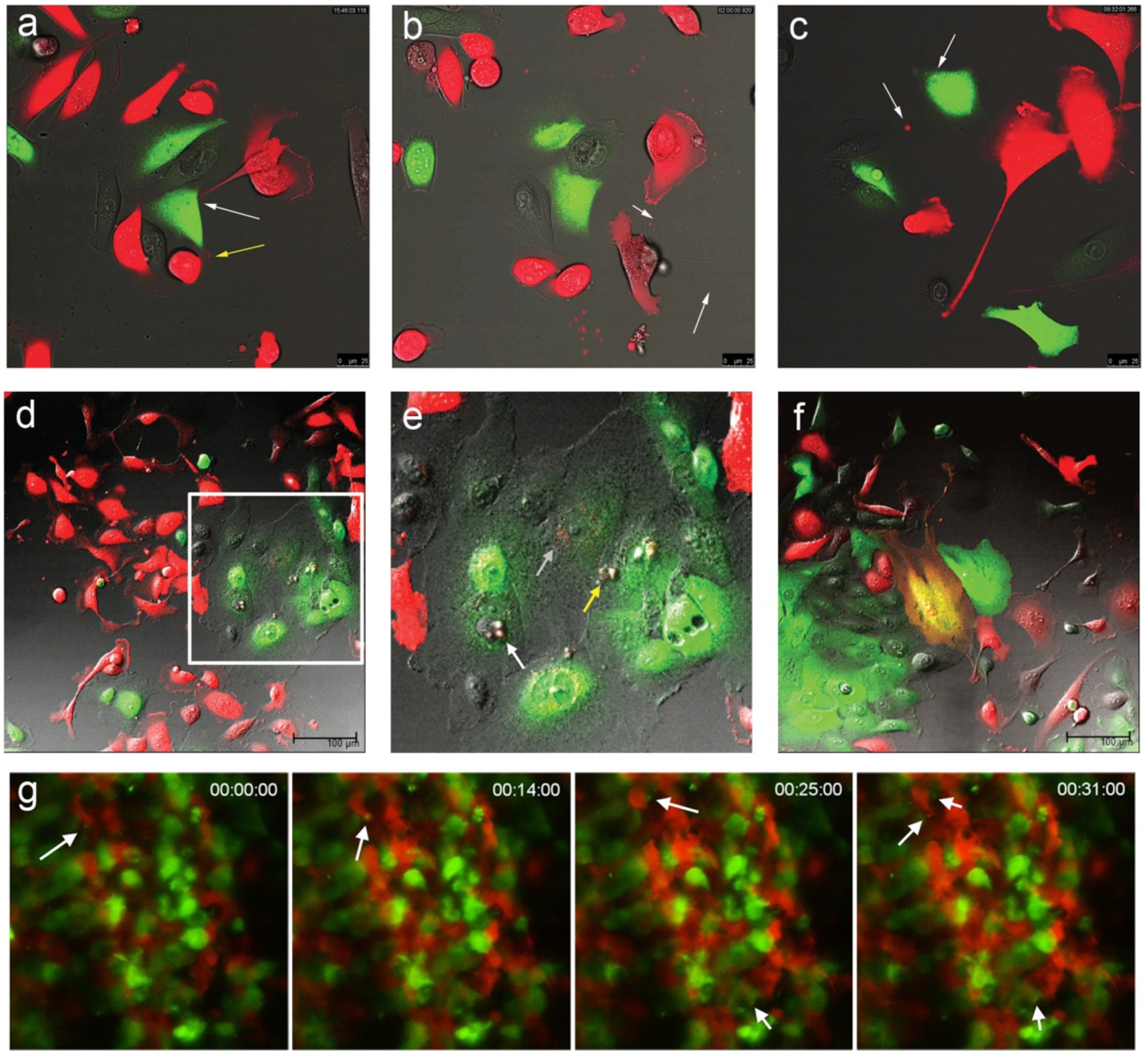
Dynamic cell-cell interactions between breast cancer cells and non-tumorigenic mammary epithelial cells. Representative images of cell-cell interactions between RFP-transfected MDA-MB-231 breast cancer cells and GFP-transfected MCF10A non-tumorigenic mammary epithelial cells were captured using time-lapse confocal microscopy (a-d). **a**. MDA-MB-231 cells use lamellipodia-like structures to contacting MCF10A cells (yellow arrow) or distant MCF10A cells (white arrow). **b**. Nanotube-like structures extending from MDA-MB-231 cells are seen (white arrows). **c**. RFP-expressing vesicles from MDA-MB-231 cells are transferred to MCF10A cells (white arrows). **d-e**. The area within the white rectangle in Figure 1d is shown in 1e. Transferred vesicles from MDA-MB-231 cells are located in both nucleus (white arrow) and cytoplasm (yellow arrow) of MCF10A cells. **f**. A minority of co-cultured cells show dual fluorescence. **g**. The representative images of the *in vivo* behavior of MDA-MB-231 cells and MCF10A cells co-injected in the earlobe of mouse are shown with migrating extracellular vesicles (white arrows).

Among the various cell-cell interactions, the exchanges of extracellular vesicles were frequently observed. MCF10A cells engulfed extracellular vesicles originated from MDA-MB-231 cells, and the vesicles were often transferred to the nucleus while some stayed at cytoplasm (Figure 1d and 1e). Additionally, a small proportion of co-cultured cells (1∼2%) exhibited mixed fluorescence (Figure 1f, Supplementary Figure S2). To observe the cell-cell interactions *in vivo*, we inoculated a mixture of MDA-MB-231 cells and MCF10A cells in the earlobes of nude mouse and obtained the time-lapse imaging data. After one hour of the inoculation, the movements of the MDA-MB-231 cells around the MCF10A cells were detectable (Supplementary Video S3). After 24 hours, the cell-cell interactions and exchange of vesicles were more frequently seen between the cells (Figure 1g, Supplementary Videos S4). These data suggest that the cancer cells and epithelial cells can undergo a dynamic range of physical contacts and cell-cell interactions both *in vitro* and *in vivo*.

### Direct co-culture with breast cancer cells induce phenotypic changes in MCF10A cells

We then determined whether the cell-cell interactions with breast cancer cells induce phenotype changes in MCF10A cells. First, we co-cultured MCF10A cells with breast cancer cell lines using in-direct co-culture system. MCF10A cells co-cultured indirectly with cancer cells showed no significant changes in cell morphology or the cell growth patterns in 2D and 3D cultures (Supplementary Figure S3). However, when the MCF10A cells were cultured in the direct co-culture system with breast cancer cells, the MCF10A cells showed substantial phenotypic changes. Directly co-cultured MCF10A cells, sorted by GFP-expression, showed changes in cell morphology such as transition into spindle-shaped cells and loss of cell-cell adhesions (Figure 2a). Additionally, the MCF10A cells showed marked decrease in E-cadherin expression when they were directly co-cultured with MDA-MB-231 cells (Figure 2b).

**Figure 2.**
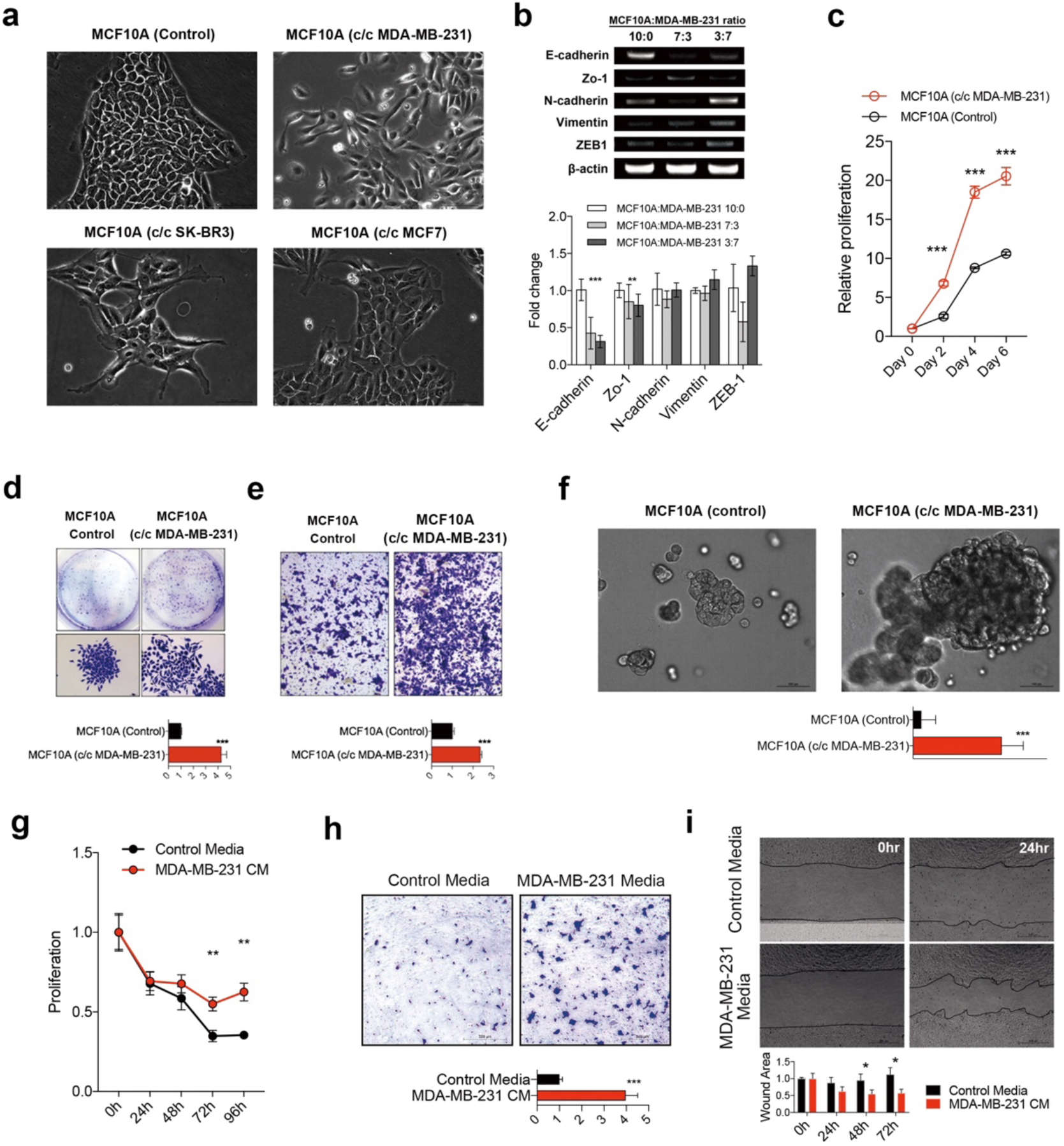
Phenotype transition of non-tumorigenic mammary epithelial cells when directly co-cultured with breast cancer cells. **a**. MCF10A cells were directly co-cultured with MDA-MB-231, SK-BR3, and MCF7 breast cancer cells. After sorting out the MCF10A cells, the cellular morphologies are shown. **b**. The expression levels of epithelial-mesenchymal transition markers were measured by PCR in MCF10A cells after co-culture with MDA-MB-231 cells (upper) and quantified (lower). **c**. Relative growth rate of co-cultured MCF10A cells was measured based on ATP level with CellTiter-Glo reagent. The representative images and quantified results of the colony-formation assay (**d**), the transwell migration assay (**e**), the 3-dimensional Matrigel culture assay (**f**) for co-cultured MCF10A cells are shown. The results of cell viability (**g**), transwell migration (**h**), and wound-healing assay (**i**) are shown after treating MCF10A cells with the conditioned media obtained from MDA-MB-231 cells. Error bars denote mean ± SD. *P < 0.05, **P < 0.01, ***P < 0.001. P values are determined by the Mann-Whitney test.

When the isolated MCF10A cells were grown in vitro, the proliferation rate, colony formation capacity, and the cell migration ability were significantly enhanced in cells directly co-cultured with MDA-MB-231 cells (Figure 2c-2e). MCF10A cells also showed higher numbers and larger sizes of cell spheres when they were grown in 3D matrigels (Figure 2f). Although the indirect co-culture with MDA-MB-231 cells did not cause significant morphologic changes in MCF10A cells, the MCF10A cells were more resistant to the serum deprivation (FIgure 2g) and showed increased migration capacity as measured by transwell migration assay and the wound-healing assay (Figure 2h-2i) when they were treated with the conditioned media from the MDA-MB-231 cells. Collectively, our data indicate that the non-tumorigenic mammary epithelial cells may undergo a substantial phenotype changes induced by breast cancer cells and the effect is more pronounced within the milieu where the non-tumorigenic mammary epithelial cells experience direct contacts with the cancer cells.

### Gene expression changes and signaling dysregulation occurring in the co-cultured MCF10A cells reflect tumor microenvironment

To understand the molecular mechanisms underlying the phenotype changes induced by direct co-culture, we analyzed the transcriptome profiles of MCF10A cells when they were directly co-cultured with MDA-MB-231 cells. A total of 151 genes were dysregulated in directly co-cultured MCF10A cells more than two-fold in expression (Figure 3a, Supplementary Table S1). Again, we observed the decreased expression of mammary epithelial differentiation markers such as CDH1 or CD24^17^. Based on the gene expression profiles, we identified several cancer-related pathways, including metabolic pathway, cell adhesion, growth factor signaling, and TP53 pathway, that were significantly dysregulated in MCF10A cells after the direct co-culture (Figure 3b).

**Figure 3.**
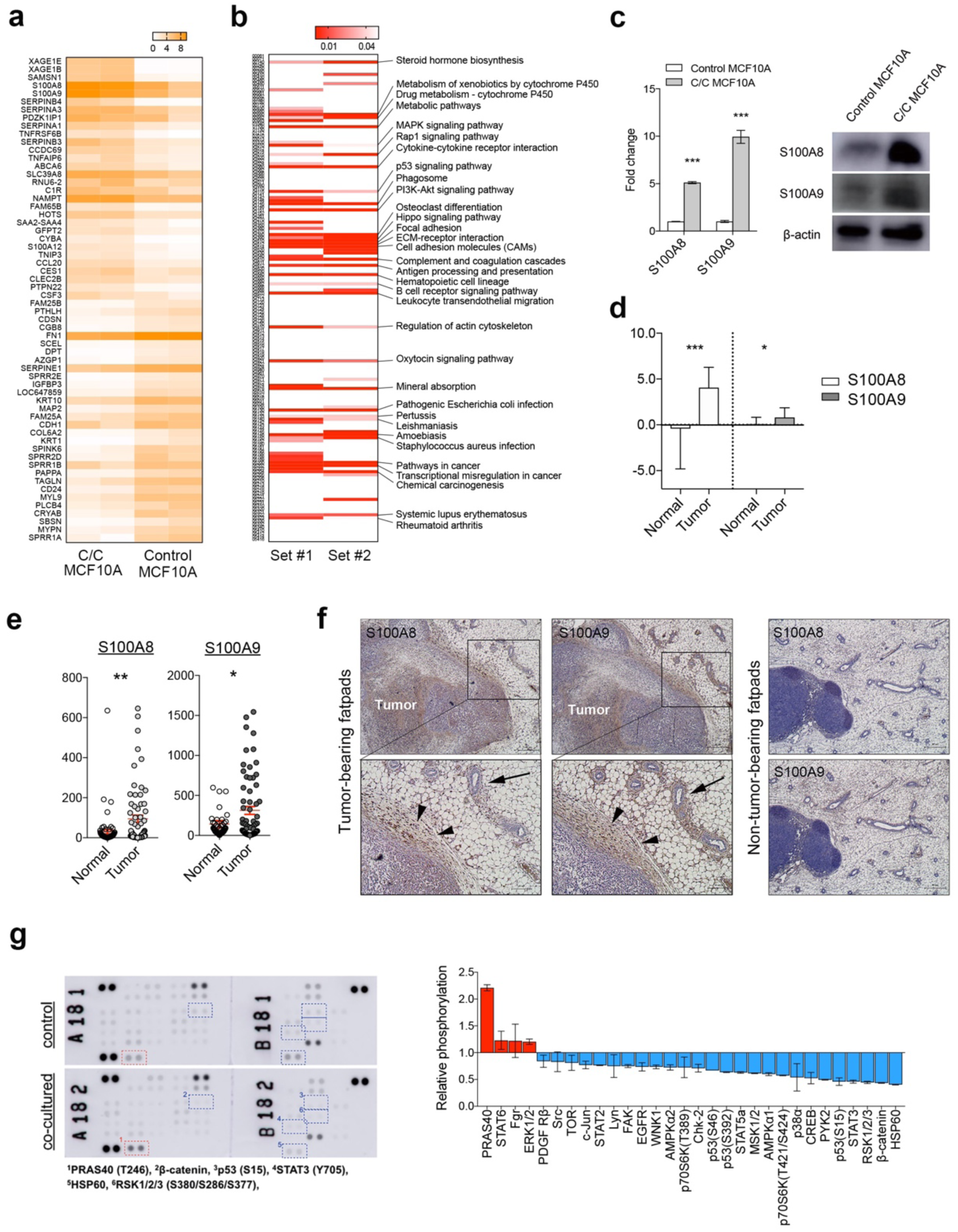
Transcriptomic and proteomic profiles of non-tumorigenic mammary epithelial cells co-cultured with breast cancer cells. **a**. Heatmap shows the top 60 genes that were up- or down-regulated in MCF10A cells co-cultured with MDA-MB-231 cells. **b**. Heatmap shows the significant enrichment of KEGG pathways based on the RNA sequencing data. Pathways significantly dysregulated in both datasets are listed on the right side. **c**. S100A8/A9 mRNA levels and protein levels in MCF10A cells co-cultured with MDA-MB-231 cells are shown. **d**. S100A8/A9 mRNA levels in breast cancer stromal tissues are shown (Finak et al. GSE9014). **e**. The S100A8/A9 mRNA levels obtained from human breast cancer RNA sequencing data are shown. **f**. On left, mouse mammary fatpad tissues bearing 4T1 breast cancer cells were stained against S100A8 and S100A9. Lower panels show magnified images of the upper insets. S100A8 and S100A9 proteins were detected in peri-tumoral stromal tissues (black arrowheads) and adjacent normal epithelial cells (black arrows). On right, non-tumor-bearing mouse fatpad were stained with S100A8 and S100A9. **g**. Results of phospho-protein array experiments are shown. Spots with more than two-fold changes are marked with numbered rectangles and listed below (upper). The overall quantification results of phospho-protein array are shown (lower). *P < 0.05, **P < 0.01, ***P < 0.001. P values are determined by the unpaired Student t-test for d and e and Mann-Whitney test for c. C/C denotes for co-cultured MCF10A cells.

Among the differentially expressed genes, S100A8 and S100A9 genes were highly upregulated in directly co-cultured MCF10A cells and their increased expression was validated in mRNA and protein levels (Figure 3c). Additionally, the mRNA dataset from Finak et al^18^ showed that S100A8 and S100A9 genes were significantly upregulated in the adjacent stroma tissues in human breast cancer (Figure 3d). Furthermore, S100A8 and S100A9 mRNA levels in normal breast and breast cancer tissues were examined using RNA sequencing data comprised of 65 normal and 68 cancer tissues, both genes were significantly upregulated in breast cancer tissues when compared to normal breast tissues (Figure 3e).

We then used mouse mammary carcinoma xenograft model to determine whether the induction of S100A8/A9 expression by breast cancer cells also occurs *in vivo*. BALB/c mouse fatpads were injected with 4T1 murine mammary carcinoma cells and were subsequently harvested. Compared to non-tumor-bearing fatpads, mouse mammary epithelial cells and peritumoral stromal cells showed increased expression of both S100A8 and S100A9 (Figure 3f). These findings in addition to the above transcriptome data indicate that breast cancer cells induce S100A8/9 expression in tumor microenvironment especially in non-tumorigenic mammary epithelial cells.

To determine the changes in the cell signaling in co-cultured MCF10A cells, we used the phosphorylation antibody array to profile the phospho-protein signaling pathways. Many of the signaling pathways showed substantial dysregulation. PRAS40, STAT6, ERK1/2 levels were substantially increased while HSP60, β-catenin, RSK1/2/3, STAT3, and p53 levels were downregulated (Figure 3g). These results suggest that, in addition to the changes in the gene expression profiles shown above, non-tumorigenic mammary epithelial cells undergo a significant shift in signaling pathways when co-cultured with cancer cells.

### S100A8/A9 upregulation contribute to the phenotypic and molecular changes in the co-cultured MCF10A cells

To determine the functional importance of S100A8 and S100A9 gene expression in mammary epithelial cells, we established a stable MCF10A cells that overexpress S100A8 and S100A9 genes. Transduction of S100A8 overexpression vector resulted in upregulation of both S100A8 and S100A9 proteins (Supplementary Figure S4). S100A8-overexpressing MCF10A cells showed higher rate of cell proliferation when compared to the control cells (Figure 4a). Furthermore, S100A8-overexpressing MCF10A cells showed increased cell migration, invasion, colony formation, and 3-dimensional cell growth which are characteristics of the MCF10A cells directly co-cultured with breast cancer cells. (Figure 4b-4d).

**Figure 4.**
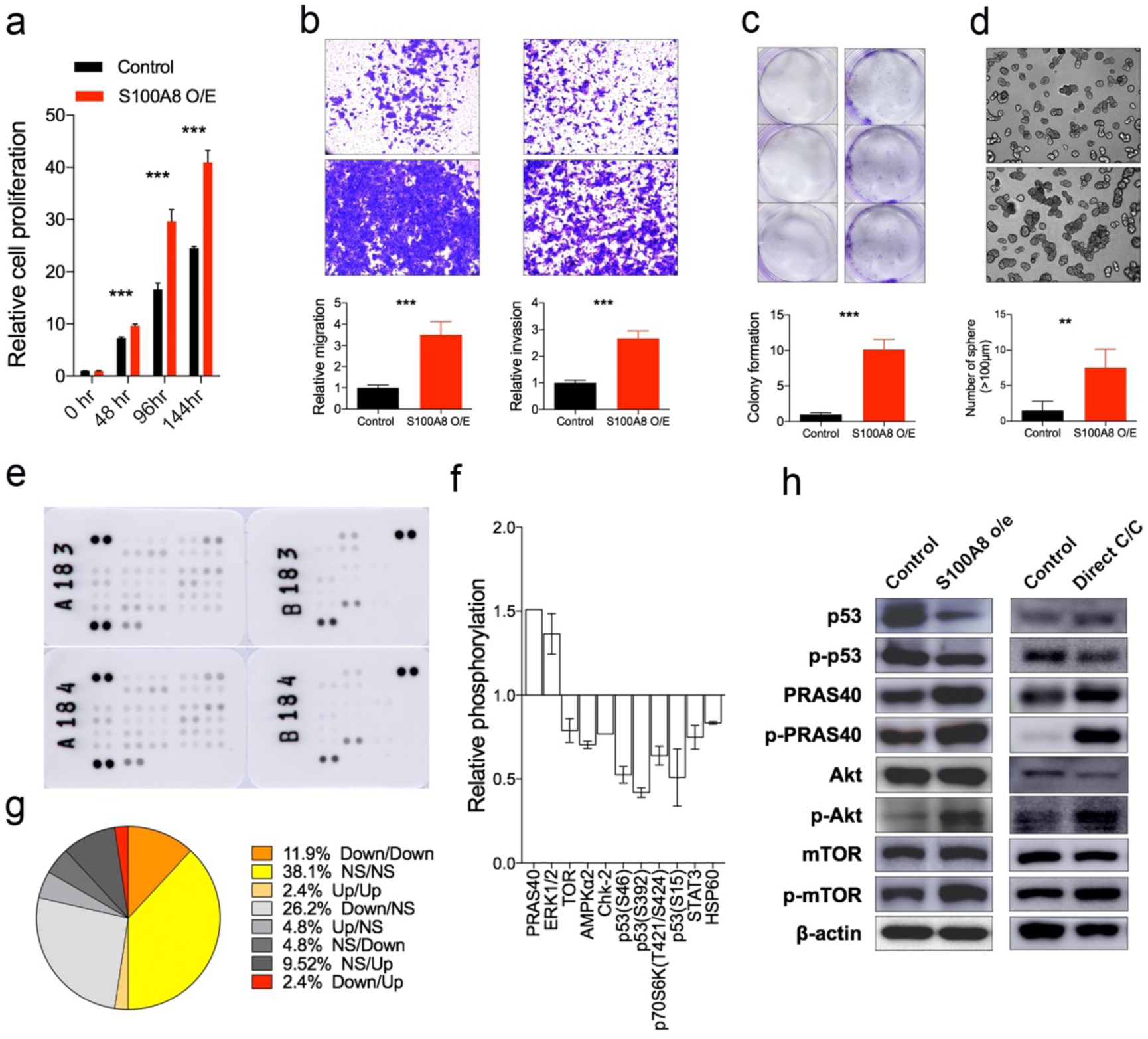
S100A8/A9-overexpression induce the phenotypes of non-tumorigenic mammary epithelial cells co-cultured with breast cancer cells. **a**. Relative growth rate of S100A8/A9-overexpressing MCF10A cells was determined based on ATP level with CellTiter-Glo reagent. **b**. The representative images and quantified results of transwell migration assay for S100A8/A9-overexpressing MCF10A cells are shown. **c**. The representative images and quantified results of Matrigel invasion assay for S100A8/A9-overexpressing MCF10A cells are shown. **d**. The representative images and quantified results of colony-formation assay for S100A8/A9-overexpressing MCF10A cells are shown. **e-f**. Results of phosphor-protein array experiments with S100A8/A9-overexpressing MCF10A cells (e) and quantified expression of significantly dysregulated proteins (f) are shown. **g**. The comparison of phospho-protein array patterns for co-cultured MCF10A cells and S100A8/9-overexpressing MCF10A cells are shown. For example, ‘Down/Up’ designates spots that are down-regulated in co-cultured MCF10A cells and up-regulated in S100A8/A9-overexpressing MCF10A cells, respectively. **h**. Western blotting results showing the protein expression levels of selected dysregulated proteins in both co-cultured MCF10A cells and S100A8/A9-overexpressing MCF10A cells. Error bars denote mean ± SD. ^**^P < 0.01, ***P < 0.001. P values are determined by the Mann-Whitney test.

To investigated the effect of S100A8 and S100A9 expression on the signaling pathway activities, we repeated the phosphorylation antibody array in S100A8-overexpressed MCF10A cells (Figure 4e-4f). More than half of the signaling proteins observed in S100A8-overexpressed MCF10A cells showed concordant expression patterns with directly co-cultured MCF10A cells and only one protein showed opposite expression pattern between the two cells (Figure 4g). The dysregulation of signaling pathways in both S100A8 overexpressing MCF10A cells and the MCF10A cells directly co-cultured with MDA-MB-231 cells were further validated using western blotting as shown in Figure 4h. Taken together, out data demonstrate that, at least in part, the phenotypic and molecular transitions seen in the mammary epithelial cells when co-cultured with breast cancer cells can be explained by the upregulation of S100A8/A9.

## Discussion

Normal mammary epithelial cells are closest neighboring cells to malignant epithelial cells during the initial process of carcinogenesis. Also, cancer cells often contact adjacent normal epithelial cells during the invasion process. The underlying hypothesis of our work was that normal epithelial cells adjacent to cancer cells may undergo molecular and functional transition despite the normal microscopic appearance. Indeed, we observed that non-tumorigenic epithelial cells undergo a substantial molecular and phenotypic changes when the cells were directly exposed to the breast cancer cells. Non-tumorigenic mammary epithelial MCF10A cells showed significant increase in cell proliferation and migration when they were directly co-cultured with breast cancer cells. Along with the changes in cell behavior, directly co-cultured MCF10A cells also exhibited spindle-shaped morphologies and significant downregulation of epithelial adhesion markers such as E-cadherin and Zo-1. Previous studies have shown that E-cadherin expression in normal epithelial cells plays an important role during the process initial carcinogenesis by regulating cell protrusion formation of neighboring transformed cells.^12,19^ Additionally, three-dimensional direct co-culture with breast cancer cells induced loss of epithelial differentiation features in non-tumorigenic MDCK cells.^20^ Finally, Trujillo et al^21^ have shown that normal epithelial cells adjacent breast tumors show increased expression of EMT markers such as α-SMA and S100A4. Our data, along with these previous reports, suggests that breast cancer cells may induce epithelial mesenchymal transition in adjacent normal mammary epithelial cells via direct cell-cell contacts.

Using the *in vitro* direct co-culture approach, we observed dynamic and complex cell-cell interactions between breast cancer cells and non-tumorigenic mammary epithelial cells that include diverse physical contacts and vesicle transfers. We were also able to demonstrate that the tumor cells and normal epithelial cells exhibit various physical interactions *in vivo* by showing live images taken from the mouse earlobe model. Such dynamic interactions have also been reported in a three-dimensional co-culture experiment^21^ and between various stromal cell types in microenvironment^22^. Recently, Roh-Johnson et al^23^ have shown that the cytoplasmic transfer during the cell-cell contacts between melanoma cells and macrophages is critical in cancer cell invasion *in vivo*. The clinical and biologic consequences of this cell-cell interactions between malignant cancer cells and adjacent normal epithelial cells are unclear. Normal epithelial cells exert tumor-suppressive effects during the initial phase of carcinogenesis by cell competition and protrusion of neighboring transformed cells.^19,24,25^ Alternatively, tumor cells can reprogram the adjacent normal epithelial cells to form a pro-tumorigenic microenvironment at the late stages of carcinogenesis.^13^ This phenomenon of tumor-driven reprograming of microenvironment has been shown for other cell types such as fibroblast or adipocytes in breast cancer.^5,6^ Further research is needed to clarify the role of molecular transformation of normal epithelial cells during the breast cancer progression.

In molecular levels, we observed that S100A8/A9 expression levels are significantly up-regulated when non-tumorigenic MCF10A epithelial cells were co-cultured with MDA-MB-231 cells. The up-regulation of S100A8/A9 was also observed in a syngeneic mouse xenograft model of 4T1 murine mammary carcinoma cells. S100A8/A9 act as alarmins that trigger damage-associated molecular patterns molecules and modulate the immune response.^26^ S100A8/A9 overexpression in MCF10A cells resulted in a similar cell phenotype seen in directly co-cultured MCF10A cells in terms of in vitro cell behaviors and cell signaling activities. Moon and colleagues have previously shown that S100A8/A9 overexpression can induce an invasive signaling program in breast epithelial cells via the H-RAS activity.27,28 Our results show that S100A8/A9, potentially critical regulators of cell behavior, could be induced in non-tumorigenic mammary epithelial cells during the dynamic cell-cell interactions with adjacent breast cancer cells. Additionally, the directly co-cultured and S100A8/A9-overexpressing cells both showed increased phosphorylation of PRAS40-Thr^246^ and its upstream Akt-Ser^473^. Phosphorylation of Akt-Ser^473^ and PRAS40-Thr^246^ activate mTOR signaling that affects diverse biologic aspects of the epithelial cells and other cells of tumor microenvironment.^29-31^ The biologic implications and therapeutic potentials of the observed mTOR signaling dysregulation in mammary epithelial cells in breast cancer microenvironment should be further explored. Our study has several limitations. First, we have used a limited number of cell lines during the present study. The molecular transitions in non-tumorigenic mammary epithelial cells might vary along the different subtypes of breast cancers. Second, the effect of this phenotype changes in mammary epithelial cells on the breast cancer cells in terms of breast cancer growth and metastasis has not been investigated using *in vivo* breast cancer models. Finally, although we’ve demonstrated the up-regulation of S100A8/A9 expression in mouse fatpad xenograft tumors, we were not able to examine the S100A8/A9 up-regulation in human breast cancer tissues. Further studies investigating the biologic consequence and clinical implications of the molecular changes in the non-tumorigenic mammary epithelial cells in breast cancer microenvironment are needed.

In conclusion, our study demonstrate that breast cancer cells may induce substantial molecular changes in non-tumorigenic mammary epithelial cells via dynamic cell-cell interactions. As the results, mammary epithelial cells undergo a phenotype transition which involves more active proliferation and migration. S100A8/A9 may play a pivotal role during this phenotype transition of mammary epithelial cells. Our study provides scientific basis for pursuing a novel therapeutic strategy that targets the non-tumorigenic mammary epithelial cells in tumor microenvironments.

## Methods

### Cell culture

Cells were purchased from Korean Cell Line Bank (Seoul, Korea). Non-tumorigenic mammary epithelial MCF10A cells were maintained in a 1:1 mixture of Dulbecco’s Modified Eagle’s Medium (DMEM) and Ham’s F12 medium (F12) with 5% horse serum, 20 ng/mL epidermal growth factor (EGF), 100 ng/mL cholera toxin, 10 μg/mL insulin, and 500 ng/mL hydrocortisone. MCF7 and MDA-MB-231 breast cancer cells were cultured in DMEM with 10% FBS, 1% penicillin, and 1% streptomycin. SK-BR3 breast cancer cells were cultured in RPMI 1640 with 10% FBS, 1% penicillin, and 1% streptomycin. For fluorescence tagged cells, puromycin was added.

### Co-culture of MCF10A cells and breast cancer cells

To optimize the culture medium for co-culture, we tested the effect of various mixture of MDA-MB-231 culture media and MCF10A culture media on cell survival. Based on the effect on cell proliferation, we chose 7:3 ratio mixture of MDA-MB-231 media and MCF10A media for the direct co-culture of the cells (Supplementary Figure S5). For effective separation of MCF10A cells and MDA-MB-231 cells after the direct co-culture, MCF10A cells and MDA-MB-231 cells were transfected with GFP and RFP using lentiviral vectors, respectively. After the direct co-culture, cells were isolated using a FACS Aria II cell sorter (Becton Dickinson, NJ) as shown in the Supplementary Figure S2.

Indirect co-culture was performed by using transwell inserts (pore size 0.4um). Cancer cells were seeded on the membrane of the insert and MCF10A cells were seeded on the bottom six well plates. For controls, same cells were seeded in the bottom wells and on the transwell inserts.

### *In vitro* assays measuring cell phenotypes

Proliferation assays were conducted using CellTiter-Glo Luminescent Cell Viability Assay (Promega, Madison, USA) following the manufacturer’s protocol. Cells were seeded in triplicate into 96-well plates at a density of 2,000 cells per well. For migration assay, 2×10^4^ cells were seeded in an insert (8μm pore size) with serum free media and media with 10% FBS was added in lower chambers. Cells were incubated for 20hrs and fixed with 4% paraformaldehyde and stained with 0.1% crystal violet. For colony formation assay, 2×10^3^ cells were seeded into 6-well plates. After 2 weeks, colonies were fixed in 4% paraformaldehyde and stained with 0.1% crystal violet.

### RNA Sequencing and qPCR

RNA sequencing libraries were prepared using TruSeq RNA Access library kit (Illumina, Inc., San Diego, CA, USA) according to the manufacturer’s protocol. After validation of the libraries, using Agilent DNA screentape D1000 kit on a TapeStation (Agilent Technologies, Santa Clara, CA, USA), the hybridization steps were performed using exome capture probes and streptavidin coated beads. RNA sequencing was performed by HiSeq 2000 (Illumina,San Diego, USA by the Macrogen Incorporated). We processed reads from the sequencer and aligned them to the Homo sapiens (hg19). After aligning reads to genome, Cufflinks v2.2.1 was used to assemble aligned reads into transcripts and to estimate their abundance. The transcript counts in isoform and gene level were calculated, and the relative transcript abundances were measured in FPKM (Fragments Per Kilobase of exon per Million fragments mapped) from Cufflinks.

RNA sequencing data of breast cancer was generated through a separate project to characterize the genomic profiles of the primary breast tumor and patient-derived xenograft tumors which will be presented in other reports (IRB No. 1402-054-555). For qPCR, total RNA was extracted from isolated cells with TRIzol (Favorgen, Taiwan). PrimeScript 1st strand cDNA Synthesis Kit (Takara, Japan) were used for reverse transcription of RNA, and resulting cDNA was amplified using Power SYBR® Green PCR Master Mix (Applied Biosystems, CA). The information on the primer sequences used in this study is listed in the Supplementary Table S2.

### Phosphorylation array and western blotting

For phosphorylation array, protein extraction was done with buffer following the manufacturer’s protocol. Protein concentration was measured by BCA assay kit (Thermo scientific, Palm Springs, CA, USA). Phosphorylation array was performed by using the human phosphor-kinase array kit (R&D systems, USA).

Proteins were harvested with RIPA buffer (Thermo scientific, Palm Springs, CA, USA), protease & phosphatase inhibitor and 0.5M EDTA solution. Protein concentration was measured by BCA assay kit (Thermo scientific, Palm Springs, CA, USA). Cell lysates were loaded onto 10% gels and transferred to a PVDF membrane. The membrane was blocked with 5% skim milk and incubated with primary antibody overnight at 4°C. The peroxidase-conjugated secondary antibody was used for detection. Bands were detected by LAS. The information on the antibodies used in this study is listed in the Supplementary Table S3.

### Cell imaging

For time-lapse imaging of cells during the direct co-culture, 9×10^3^ GFP-transfected MCF10A cells and 2×10^4^ RFP-transfected MDA-MB-231 cells were seeded in 8-well chambers and cultured for 24 hours. After 24 hours culture, live cells images were taken every 10 minutes with fixed position for 16 hours by using Leica confocal microscope (Leica TCS SP8, Leica Microsystems Ltd, Korea), and the obtained images were analyzed by Leica LAS X program.

For *in vivo* imaging, the experiments were performed in accordance with the Animal Care and Use Committee guidelines of Woosung BSC (Suwon, Korea). Hair-removed ear skin of C57BL/6 mouse (10 weeks old) was injected with 4.6 × 10^4^ cells of the mixture MCF10A-GFP (2.3 × 10^4^) and MDA-MB-231-RFP (2.3 × 10^4^) cell lines. The mice were anesthetized with an intraperitoneal injection of Zoletil (30 mg/kg, Virbac, Carros, France) and Rompun (10 mg/kg, Bayer-Korea, Seoul, Korea) before imaging. The mice were placed on the heated plate of a motorized XYZ translational stage. The *in vivo* movement of MCF10A and MDA-MB-231 was monitored by modified custom-built laser-scanning confocal microscopy {Choe, 2015 #3}. GFP-expressing cells were visualized at an excitation wavelength of 491 nm and detected through a bandpass filter of 502 nm to 537 nm (Semrock Inc, Rochester, NY, USA). RFP-expressing cells were imaged at an excitation wavelength of 532 nm and detected using a bandpass filter of 562 nm to 596 nm (Semrock Inc). Cell movement was visualized at 1 min interval for 2 hr. After acquired from the imaging system, 512 × 512 pixel images were then compensated with Matlab (Mathworks, Natick, MA, USA) and reconstructed by ImageJ software.

### S100A8/A9 expression in xenograft tumor-bearing mouse fatpad

5-week-old BALB/c mice were purchased from KOATECH (Seoul, Korea) and housed in Seoul National University Hospital Clinical Research Institute’s Specific Pathogen Free zone. All experiments were approved by the Institutional Animal Care and Use Committee in Seoul National University Hospital (SNUH-IACUC, 17-0165-S1A0 (1)). To obtain control mouse fatpads, eight-week-old non-tumor-bearing BALB/c mouse was sacrificed and mammary fat pad was resected. To obtain tumor-bearing mouse fat pad, 2 × 10^5^ 4T1 mouse mammary epithelial cancer cells were injected in 6-week-old BALB/c mouse’s mammary fat pad. At two weeks after the tumor injection, the mouse was sacrificed and tumor-bearing mammary fatpad was resected. The resected fatpads were fixed with 4% PFA and embedded in paraffin.

### S100A8/A9-overexpressing MCF10A cell line

The coding sequences of S100A8 and S100A9 were acquired by RT-PCR and cloned into pCDH-GFP and pCDH-RFP vectors. MCF10A cells were transfected with lentivirus S100A8-GFP and S100A9-RFP construct. Transfected cells were selected by puromycin.

## Statistical analysis

In general, most data represent the mean ± S.D and are representative of 3 independent experiments, except for RNA sequencing and phospho-protein arrays which are two independent experiments. Graph Pad Prism (ver. 7.01) was used for generating graphs and heatmaps and performing statistical tests. P values were calculated from unpaired two-tailed Student’s t tests or Mann-Whitney test as appropriate.

To identify differentially expressed genes, we filtered genes with one more than zeroed FPKM values and the data were log2-transformed and subjected to quantile normalization. Statistical significance of the differential expression data was determined using fold change in which the null hypothesis was that no difference exists among samples. Gene pathway analysis for the DEG was done based on KEGG pathway (http://www.genome.jp/kegg/pathway.html).

## Data availability

All data supporting our findings can be found in the main paper or in supplementary files.

## Author contributions

SHJ performed the bulk of the work reported, including writing the initial draft, generating cell lines, various in vitro studies, and phospho-protein arrays. WHH and MQ performed fatpad tumor xenograft studies, PCR, and western blotting. BSH, JHK performed YP and DSL assisted and provide guidance to cell sorting and animal experiments. NHK and MCP performed in vivo cell imaging studies. HBL, and WH analyzed the transcriptome data and pathway analysis. JC and JIK provided RNA sequencing data of breast cancer and analyzed the target gene expression levels. DYN and HGM initiated the study, provided overall direction and oversight to the project, and finalized the manuscript. All authors read and approved the final manuscript.

