## Supplementary Table S2-S3 for "S100A8/A9 mediate the reprograming of normal mammary epithelial cells induced by dynamic cell-cell interactions with adjacent breast cancer cells"

Supplementary Table S2. Information of primers used in this study.

| Name | | Sequences |
|---|---|---|
| S100A8 | F | ATGCCGTCTACAGGGATGAC |
| | R | CCACGCCCATCTTTATCACC |
| S100A9 | F | GGGAATTCAAAGAGCTGGTGC |
| | R | AGCTGCTTGTCTGCATTTGTG |
| KRT17 | F | CAGTCCCAGCTCAGCATGAA |
| | R | CCACAATGGTACGCACCTGA |
| KRT16P3 | F | TCAGACCGGTGGAGAAGTGA |
| | R | TGTGCCGGGTCCTTCATACT |
| KRT16 | F | AGTCCCAGCTCAGCATGAAA |
| | R | GCGGGAAGAATAGGATTGGC |
| KRT10 | F | CAACTCACATCAGGGGGAGC |
| | R | CAGCTCATCCAGCACCCTAC |
| CD24 | F | GCTCCTACCCACGCAGATTT |
| | R | GAGACCACGAAGAGACTGGC |
| ZEB1 | F | GTGACGCAGTCTGGGTGTAA |
| | R | TGAGTCCTGTTCTTGGTCGC |
| Snail | F | GAGGACAGTGGGAAAGGCTC |
| | R | TGGTTCGGATGTGCATCTT |
| ZO-1 | F | GACAGCAGACCACGTTACGA |
| | R | TGAAGGTATCAGCGGAGGGA |
| β-actin | F | CACTGTGTTGGCGTACAGGT |
| | R | TCATCACCATTGGCAATGAG |
| E-cadherin | F | CCCTCGACACCCGATTCAAA |
| | R | TGGATTCCAGAAACGGAGGC |

Supplementary Table S3. Information of antibodies used in this study.

| Antibody | Source | Company (catalog) |
|---|---|---|
| Calgranulin A | Mouse monoclonal antibody | Santa Cruz (sc-48352) |
| Calgranulin B | Mouse monoclonal antibody | Santa Cruz (sc-376772) |
| AKT | Rabbit polyclonal antibody | Cell signaling (9272) |
| p-AKT (Ser473) | Rabbit polyclonal antibody | Cell signaling (9271) |
| p53 | Mouse monoclonal antibody | Cell signaling (2524) |
| p-p53 (Ser15) | Rabbit polyclonal antibody | Cell signaling (9284) |
| PRAS40 | Rabbit monoclonal antibody | Cell signaling (2691) |
| p-PRAS40 (Thr246) | Rabbit monoclonal antibody | Cell signaling (2997) |
| mTOR | Rabbit monoclonal antibody | Cell signaling (2983) |
| p-mTOR (Ser2448) | Rabbit monoclonal antibody | Cell signaling (5536) |
| RAGE | Rabbit polyclonal antibody | AbCam (ab3611) |
| TLR4 | Mouse monoclonal antibody | Santa Cruz (sc-293072) |
| β-actin | Mouse monoclonal antibody | Santa Cruz (sc-47778) |
