## Supplementary Figures S1-S5 for "S100A8/A9 mediate the reprograming of normal mammary epithelial cells induced by dynamic cell-cell interactions with adjacent breast cancer cells"

Supplementary Figure S1. Direct co-culture of MCF10A cells and MDA-MB-231 cells.

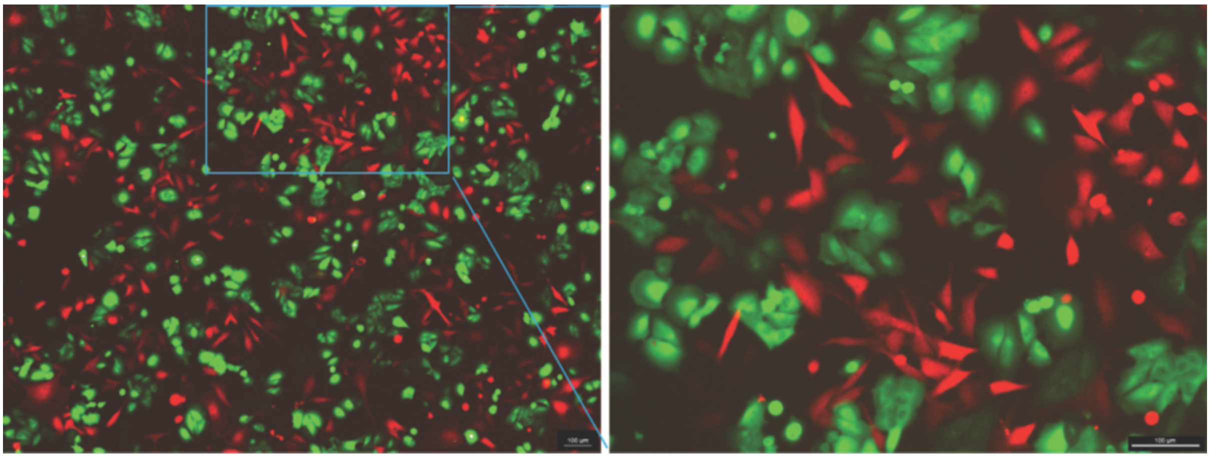

Supplementary Figure S2. Results of cell sorting after direct co-culture.

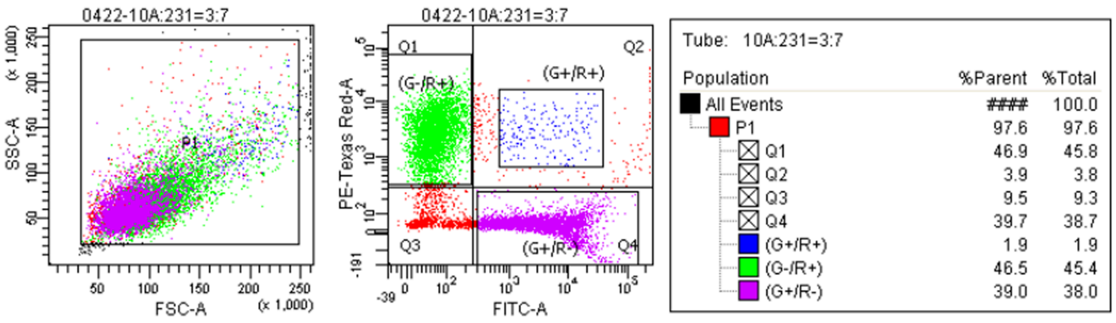

Supplementary Figure S3. MCF10A cells after indirect co-culture with breast cancer cells

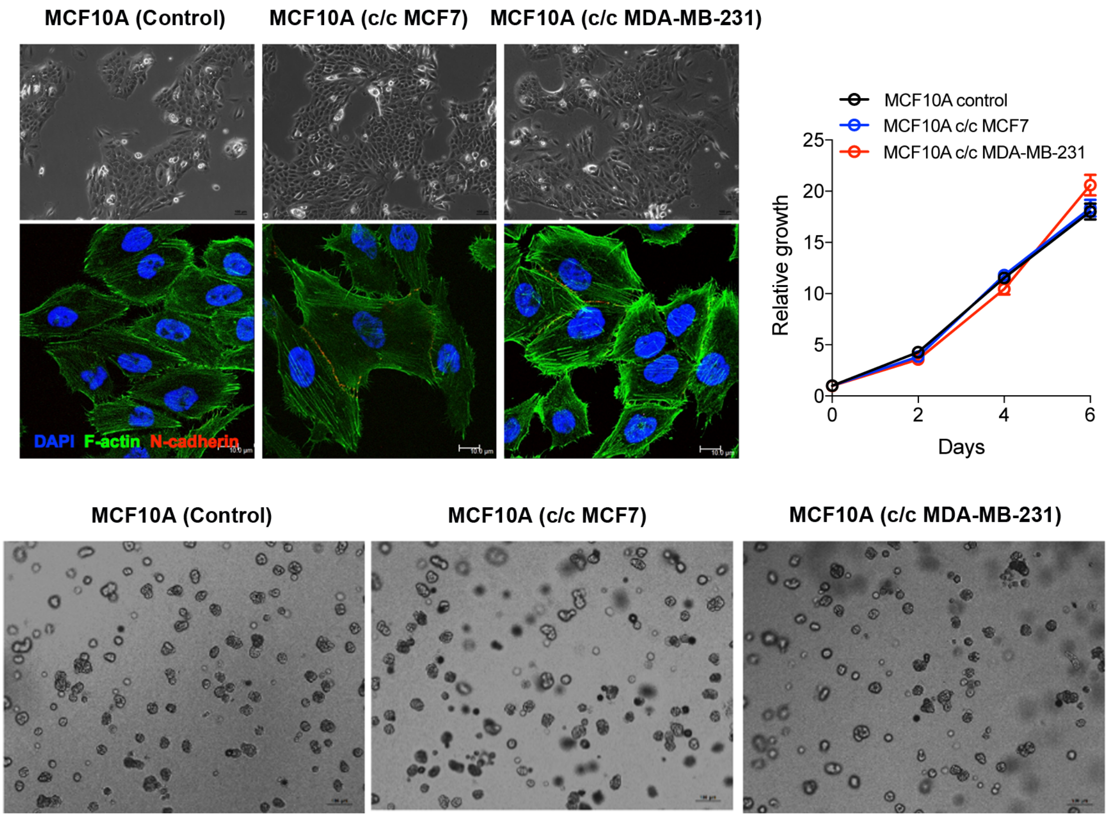

Supplementary Figure S4. Establishment of S100A8/A9-overexpressing MCF10A cells.

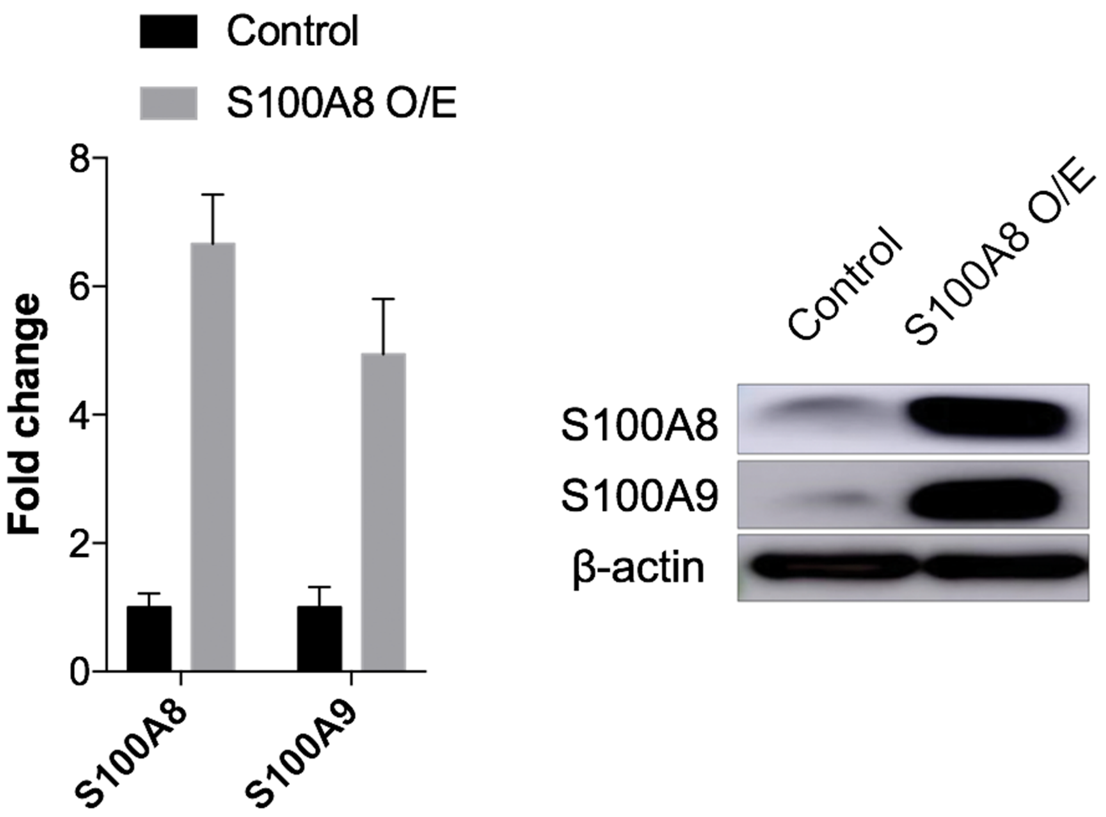

Supplementary Figure S5. *In vitro* testing of cell culture media

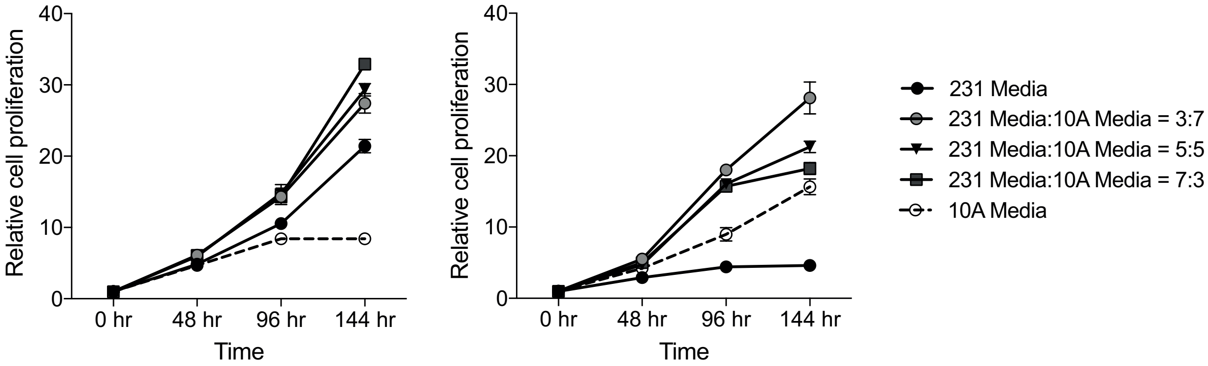
